# A stable core of GCPs 4, 5 and 6 promotes the assembly of γ-tubulin ring complexes

**DOI:** 10.1101/2020.01.21.914036

**Authors:** Laurence Haren, Dorian Farache, Laurent Emorine, Andreas Merdes

## Abstract

γ-tubulin is a major protein involved in the nucleation of microtubules in all eukaryotes. It forms two different complexes with proteins of the GCP family (gamma-tubulin complex proteins): γ-tubulin small complexes (γTuSCs), containing γ-tubulin and GCPs 2 and 3, and γ-tubulin ring complexes (γTuRCs), containing multiple γTuSCs, in addition to GCPs 4, 5, and 6. Whereas the structure and assembly properties of γTuSCs have been intensively studied, little is known about the assembly of γTuRCs, and about the specific roles of GCPs 4, 5, and 6. Here, we demonstrate that two copies of GCP4 and one copy each of GCP5 and GCP6 form a salt-resistant sub-complex within the γTuRC that assembles independently of the presence of γTuSCs. Incubation of this sub-complex with cytoplasmic extracts containing γTuSCs leads to the reconstitution of full-sized γTuRCs that are competent to nucleate microtubules. In addition, we investigate sequence extensions and insertions that are specifically found at the amino-terminus of GCP6, and between the GCP6 grip1 and grip2 motifs, and we demonstrate that these are involved in the assembly or stabilization of the γTuRC.

**Summary statement:** γ-tubulin ring complexes are templates for microtubule nucleation, composed of γ-tubulin and GCP proteins. GCPs 4, 5, 6 form a stable sub-complex, driving the assembly of the full complex.

## Introduction

γ-tubulin is a protein involved in the nucleation of microtubules. It assembles with so-called “Gamma-tubulin Complex Proteins” (GCPs) into multiprotein complexes of two different sizes. A “γ-Tubulin Small Complex” (γTuSC) comprises two molecules of γ-tubulin that are bound by GCPs 2 and 3. A much larger “γ-Tubulin Ring Complex” (γTuRC) is formed by multiple γTuSCs that associate with additional GCPs 4, 5, 6, and several smaller accessory proteins into a helical structure of 2 MDa (Kollman et al., 2011; Farache et al., 2018). A few eukaryotes, such as *Saccharomyces cerevisiae* or *Candida albicans* contain only GCPs 2 and 3. In these organisms, multiple γTuSCs are assembled that form a helix with the help of additional proteins, such as Spc110 or Mzt1 (Kollman et al., 2010; Erlemann et al., 2012; Lyon et al., 2016; Lin et al., 2014, 2016). Most eukaryotes, however, express the full complement of GCPs 2, 3, 4, 5, and 6, and form γTuRCs. It is believed that all these GCPs have similar structures. They are characterised by sequence homology in two conserved regions, the grip1 and grip2 motifs, corresponding to the N-terminal and C-terminal halves of GCP4, the smallest GCP (Gunawardane et al., 2000; Guillet et al., 2011). The crystallographic structure of GCP4 shows that these domains correspond to bundles of α-helices. The other GCPs contain additional specific sequences, mainly at the extreme N-terminus, or in the region linking the grip1 and grip2 motifs, as in GCPs 5 and 6 (Guillet et al., 2011; Farache et al., 2016).

Depletion of GCP2 or 3 leads to severe spindle abnormalities, and depleted cells are not viable. Depletion of GCPs 4, 5, or 6 can be tolerated in fission yeast or in somatic cells of *Drosophila*, but not in vertebrates, where removal of each of these GCPs prevents the formation of the γTuRC and provokes spindle defects (Anders et al., 2006; Vérollet et al., 2006; Farache et al, 2016; Cota et al., 2017). Rescue experiments with chimeric proteins containing N-terminal domains fused to C-terminal domains of a different GCP showed that the chimeras rescued the defects as long as they carried the N-terminal domain of the depleted GCP (Farache et al., 2016). Thus the GCPs are not functionally redundant, despite their structural similarities, and the function of individual GCPs are specified by their N-terminal domains. GCP2 and GCP3 interact laterally via their N-terminal domains in γTuSC helices, whereas γ-tubulin molecules are bound by the C-terminal domains (Kollman et al., 2010). FRET experiments also demonstrated a direct lateral interaction between GCP4 and GCP5 via their N-terminal domains (Farache et al., 2016), suggesting that the N-terminal domains specify lateral binding partners and thereby position the GCPs within the γTuRC helix.

In this context, the specific functions of GCPs 4, 5, and 6 in γTuRC assembly need to be investigated. Because these proteins are present only in one or two copies per complex (Murphy et al., 2001; Choi et al., 2010), and because the rescue experiments with chimeras suggest that GCPs 4, 5, 6 occupy non-random positions, their localisation within the complex is of particular interest. During the course of this work, three studies have described the structure of native γTuRCs by cry-electron microscopy, and have found a lateral association of four γTuSCs, bound to a lateral array of GCP4/GCP5/GCP4/GCP6, to which an additional γTuSC was associated (Wieczorek et al., 2020; Consolati et al., 2019; Liu et al., 2019). Altogether, this has raised the question whether GCPs 4, 5, 6 form assembly intermediates equivalent to γTuSCs. In the present study, we demonstrate biochemically that GCPs 5, 6, and two copies of GCP4 form together a stable, salt-resistant core within the γTuRC that can be purified and that drives the assembly of free γTuSCs into a full γTuRC, competent to nucleate microtubules.

## Results

### γTuRC-specific GCPs 4, 5, and 6 form a core complex resistant to high-salt-treatment

To determine how GCPs 4, 5, and 6 assemble within the γTuRC, and to examine whether γTuSC-like intermediates are formed by these proteins, we destabilised the γTuRC by treating HeLa cytoplasmic extracts with increasing concentrations of KCl. GCPs 4, 5, or 6 were immunoprecipitated and all interacting GCPs were identified by Western Blotting (Fig. 1A). Whereas the full set of GCPs was immunoprecipitated at 100mM KCl, we observed an increasing loss of γTuSCs from the immunoprecipitate at higher concentrations of KCl. At 500mM KCl, GCPs 4, 5 and 6 remained the major constituents of the immunoprecipitate, irrespective of the antibody used for precipitation. This indicated that the binding affinities between GCPs 4, 5, and 6 are stronger than their affinities to γTuSCs. Consistently, γTuRCs were previously found to dissociate and to release γTuSCs after high-salt-treatment (Moritz et al., 1998; Oegema et al., 1999).

**Figure 1:**
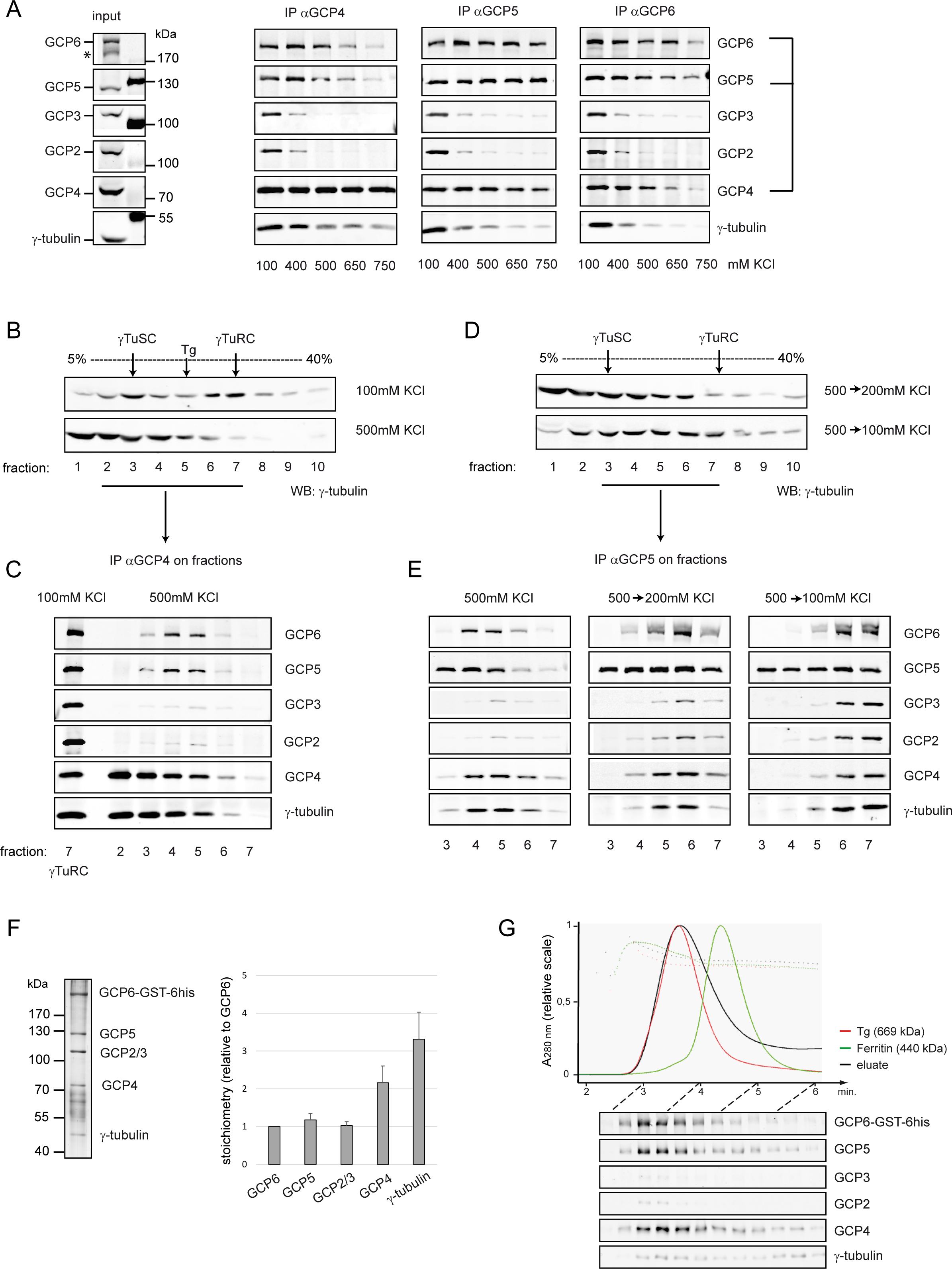
Isolation of a GCP4/5/6 sub-complex at high salt. (A) Co-immunoprecipitation of GCPs 4, 5, and 6 from HeLa cytoplasmic extract. Left panel: soluble extract (input, *: degradation product of GCP6). Panels 2, 3, 4: the extract was supplemented with increasing concentrations of KCl (100 to 750 mM), and incubated with antibodies against GCP4, GCP5, or GCP6 (IPαGCP4, IPαGCP5, IPαGCP6). Immunoprecipitated proteins were separated by SDS-PAGE, blotted and probed with antibodies against the different GCPs and γ-tubulin. Brackets on the right indicate the resistance to salt of GCPs 4, 5, 6 in the co-immunoprecipitate. (B) Sucrose gradient fractionation of HeLa cytoplasmic extract, prepared and centrifuged at 100 mM KCl (top), or adjusted to 500 mM KCl before centrifugation (bottom). Sedimentation of γTuSCs and γTuRCs was visualised by Western Blotting using an antibody against γ-tubulin. At 100 mM KCl, fraction 3 contains the majority of γTuSCs, fraction 7 the majority of γTuRCs. At 500 mM KCl, the peak of γTuRCs is lost. Thyroglobulin (“Tg”, 19.4 S) sediments in fraction 5. (C) Immunoprecipitation of GCP4 from fractions 2 to 7 of the sucrose gradient at 500 mM KCl. GCP4 co-precipitated with GCP5, GCP6, and γ-tubulin in fractions 4-5. In the first lane, the co-precipitation of all GCPs from fraction 7 of the 100 mM KCl gradient is shown as a positive control. (D) Fractionation of a HeLa cytoplasmic extract adjusted to 500 mM KCl, before centrifugation on a sucrose gradient containing 200, or 100 mM KCl (the gradient containing 500 mM KCl is shown in B, bottom row). (E) Immunoprecipitation of GCP5 from fractions 3 to 7 of the gradients containing 500, 200, or 100 mM KCl. Co-precipitation of GCP4, GCP6, and γ-tubulin was observed in peaks in fractions 4-5 at 500 mM KCl, in fraction 6 at 200 mM KCl, and in fraction 7 at 100 mM KCl. The shift of the peaks towards higher fraction numbers correlated with the co-precipitation of increasing amounts of GCP2 and GCP3. (F) Analysis of the purified GCP4/5/6 complex, from cells expressing endogenous GST-hexa-histidine-(GST-6his)-tagged GCP6, by SDS-PAGE and silver staining. The position of the core subunits was verified by Western Blot (see G). The graph on the right shows the stoichiometry of the proteins, calculated from the intensities of the bands, relative to GCP6 (mean ± SD, n=4 independent experiments). (G) Western Blot showing the fractionation of the purified GCP4/5/6 complex by gel filtration. The majority of the complex eluates with Thyroglobulin (669 kDa), and above Ferritin (440 kDa).

To investigate whether GCPs 4, 5, 6 associated in a single sub-complex within the γTuRC, we fractionated cytoplasmic extracts on gradients of 5-40% sucrose containing 100 or 500mM KCl (Fig. 1B, suppl. Fig. 1A-B). We noticed that the cell line used in these experiments (HeLa Flp-In T-REx) contains high protein levels of GCP4, part of which sedimented independently of γTuRCs, in low-density-fractions (suppl. Fig. 1A, Farache et al., 2016). To test for mutual binding of GCPs 4, 5, 6, we performed immunoprecipitation from each individual fraction in the gradient, using antibodies against GCP4 (Fig. 1C) or GCP5 (Fig. 1E, left panel). At 100mM KCl, γTuSCs and γTuRCs sedimented mainly in fractions 3 and 7, respectively, and GCPs 2 to 6 co-immunoprecipitated efficiently in fraction 7 (Fig. 1C, first lane). By contrast, at 500mM KCl we observed the disappearance of γTuRCs from fraction 7, and immunoprecipitation revealed that GCPs 4, 5, 6, and γ-tubulin associated with each other in fractions of intermediate size, excluding GCPs 2 and 3 (Fig. 1C, E, fractions 4 and 5). Interestingly, when the concentration of KCl was decreased to 200 or 100mM KCl before gradient sedimentation, the peaks of GCPs 2 to 6 shifted back to higher fractions, and GCPs 4, 5, 6 re-associated with GCPs 2 and 3 in fractions 6 and 7, respectively (Fig. 1D, E, Suppl. Fig. 1C-D). This suggests that the sub-complex of GCPs 4, 5, 6 can re-assemble with γTuSCs into γTuRCs.

We then purified the GCP4/5/6 sub-complex in order to obtain an estimation of its size and stoichiometry. We constructed a HEK293 cell line expressing GCP6 with a C-terminal GST-hexa-histidine tag, by modifying the *TUBGCP6* gene using CRISPR-Cas9. Consecutive steps of affinity-purification over glutathione sepharose and over Ni-NTA agarose were carried out in the presence of 500 mM KCl. The eluate was analysed by SDS-PAGE and silver staining, and scanned signals were normalized to the lysine molar content of each protein (Dion and Pomenti, 1983; Fig. 1F). We realized that small amounts of GCP2 or GCP3 were still present in the purified sub-complex that were probably underestimated in Western Blotting experiments (Fig. 1F). Quantifications from four independent experiments indicated a molar ratio of two copies of GCP4, one of GCP5, and one of GCP6, with three copies of γ-tubulin and one equivalent of either GCP2 or GCP3. Although GCP2 and 3 could not be distinguished by one-dimensional electrophoresis, Western Blotting proved that both proteins were present in a single band (Fig. 1F, G). Taking this stoichiometry into account, the estimated size of the complex should be around 750 kDa. Size exclusion chromatography confirmed elution at a size close to thyroglobulin, around 700kDa (Fig. 1G). These results suggest that dissociation by salt produced a heterogeneous mixture, containing a core complex made of GCP4, 5, 6, stochastically associated with either GCP2 or GCP3, and γ-tubulin.

### The core of GCP4/5/6 forms independently of γTuRC-assembly

To determine if the GCPs 4, 5 and 6 associate independently of γTuRCs, we used RNA interference (RNAi) to deplete GCP2 (Fig. 2A). Loss of GCP2 caused the disappearance of γTuRCs in sucrose gradients (Fig. 2B, suppl. Fig. 2), and GCPs 4, 5, 6 co-immunoprecipitated together with γ-tubulin in intermediate fractions. Here, the sub-complex peaked in fraction 5, whereas disassembly of γTuRCs at 500mM KCl yielded the strongest peak in fraction 4 (compare Fig. 1C to Fig. 2C). This difference may be due to the loss of interactors at high salt.

**Figure 2:**
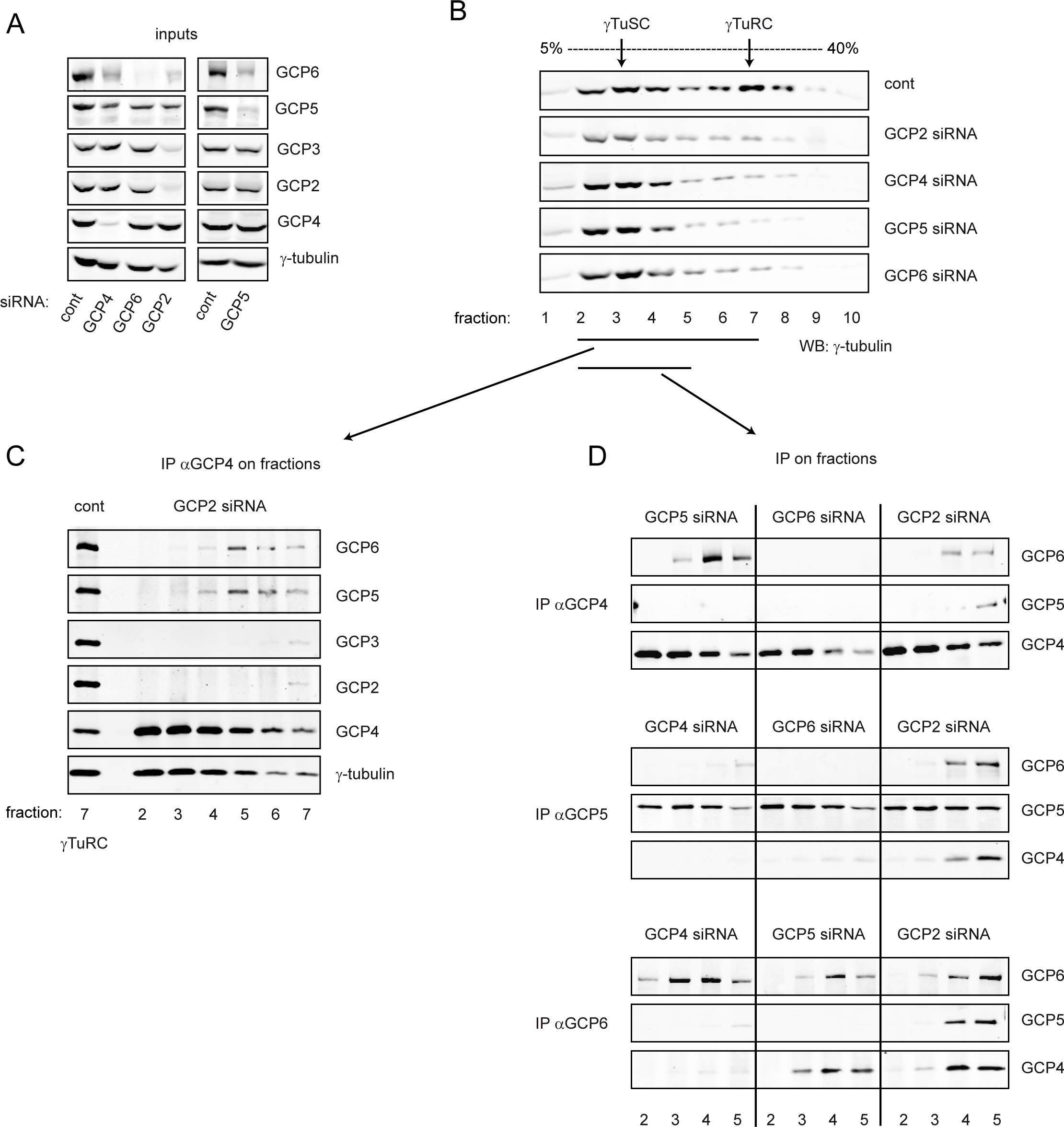
The GCP4/5/6 complex forms independently of γTuRCs. (A) Western Blot analysis of cytoplasmic extracts from control cells and cells treated with siRNAs against GCPs 2, 4, 5, or 6. The different lanes contain the inputs used in (B). Note the co-regulation of various GCPs by individual siRNAs. (B) Sucrose gradient fractionation of the extracts shown in (A), visualised with anti-γ-tubulin staining. The peak of γTuRC in fraction 7 of control extracts is lost upon siRNA-treatment against the different GCPs. (C) Immunoprecipitation of GCP4 from fractions 2 to 7, after sucrose gradient centrifugation of the cytoplasmic extract from GCP2 siRNA-treated cells. GCP4 co-precipitated with GCP5, GCP6, and γ-tubulin, mainly in fraction 5. First lane: positive control, showing co-precipitation of all GCPs from fraction 7 of a gradient from untreated cells. (D) Immunoprecipitation of GCPs 4, 5, or 6 from fractions 2 to 5, after sucrose gradient centrifugation of extracts from cells treated with siRNAs against GCPs 2, 4, 5, or 6. GCPs 4, 5 and 6 co-precipitated in the absence of GCP2, with a peak in fraction 5. GCPs 4 and 6 co-precipitated in the absence of GCP5, with a peak in fraction 4. The absence of GCPs 2 and 3 from the complexes was verified as shown in (C), but for simplicity, the corresponding Western blots were omitted in (D).

Since Wieczorek et al. (2020) and Liu et al. (2019) suggested the existence of γTuSC-like structures, containing pairs of GCPs 4/5 or GCPs 4/6, we tested how the depletion of individual GCPs 4, 5, or 6 affected the composition of the GCP4/5/6 sub-complex, and whether the proposed γTuSC-like structures did indeed exist. We depleted each of these three GCPs individually by RNAi, which resulted in each case in the loss of γTuRCs and in the accumulation of γTuSCs, as seen in sucrose gradients (Fig. 2A-B, suppl. Fig. 2). Next, we immunoprecipitated the proteins from the gradient fractions, using antibodies against either GCP4, GCP5, or GCP6 (Fig. 2D). GCPs 4, 5, and 6 were systematically co-precipitated in the absence of GCP2. GCP4 and GCP6 co-precipitated together even in the absence of GCP5. However, GCP5 bound to GCPs 4 or 6 only in the presence of all three proteins. This suggests a hierarchy of assembly, with a stable GCP4/6 small complex that enables the association with GCP5.

### Excess protein levels of GCPs 4, 5, and 6 incorporate γTuSCs into γTuRCs

We tested whether elevated amounts of GCPs 4, 5, and 6, or combinations thereof, promoted the sequestration of free γTuSCs and their incorporation into γTuRCs. On the high background of GCP4 in the HeLa Flp-In T-REx cell line (suppl. Fig. 1), GCP5 was overexpressed following transient transfection, whereas GCP6 overexpression was induced by adding doxycycline to stably transfected cells (Fig. 3A). Thus, overexpression of GCP5 resulted in an excess of both GCPs 4 and 5, but this didn’t alter the ratio between γTuSCs and γTuRCs (Fig. 3B-C). By contrast, overexpression of GCP6 resulted in an excess of GCPs 4 and 6 and led to reduced peaks of γTuSCs (fraction 3), and a simultaneous shift of protein signals of GCPs 2, 3 and γ-tubulin towards higher density fractions (Fig. 3B-D, fractions 4-7).

**Figure 3:**
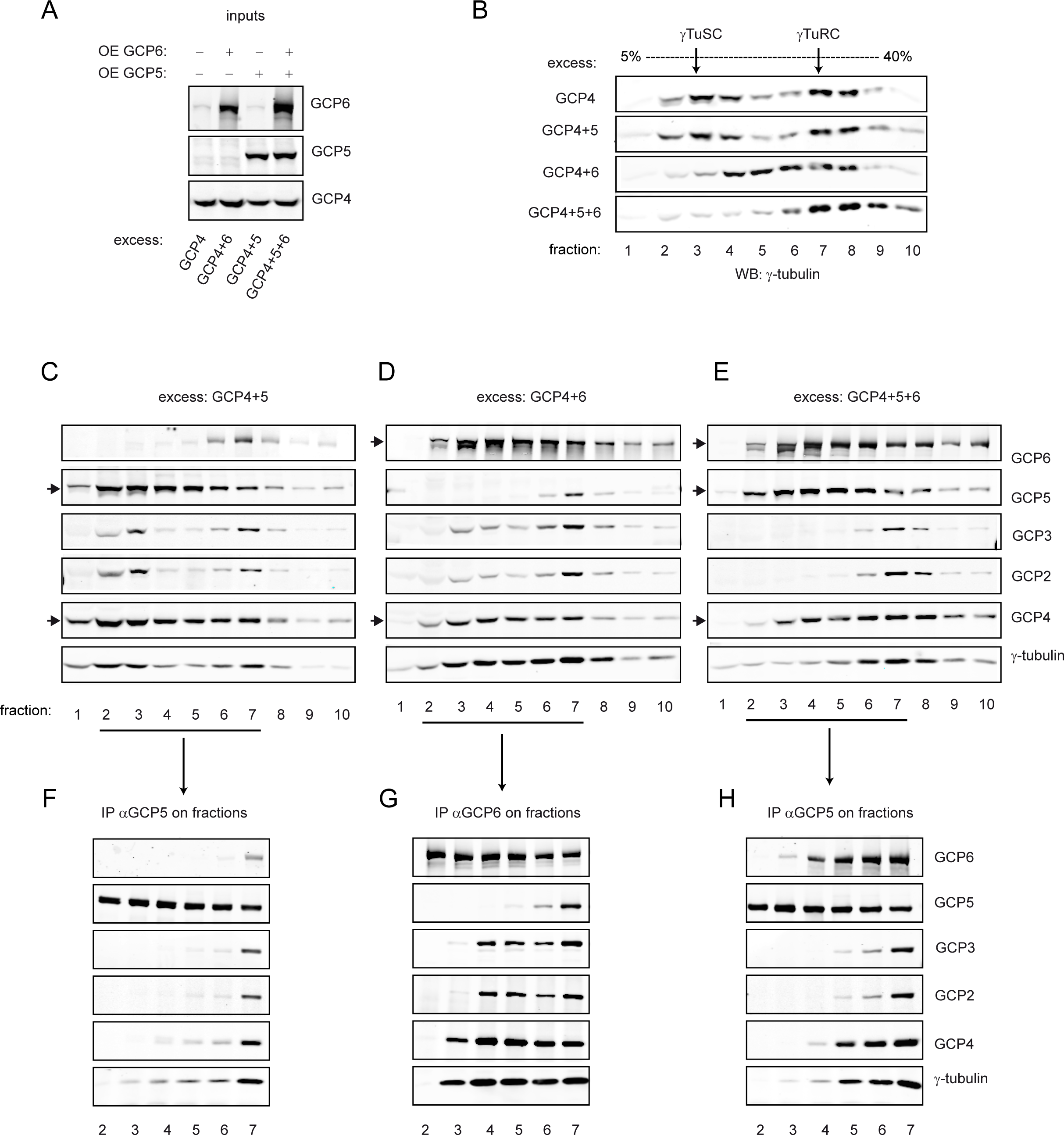
Overexpression of GCPs 4, 5, and 6 drives incorporation of free γTuSCs into γTuRCs. (A) Western Blot analysis of cytoplasmic extracts from cells with excessive amounts of GCPs 4, 5, and/or 6 (inputs for B-E). GCP4 is in excess in the cell line (see Suppl. Fig. 1). Excess of GCP5 and GCP6 was obtained by inducing the overexpression of the proteins in transiently transfected cells (“OE GCP5”), or in stably transfected cells (“OE GCP6”). (B) Sucrose gradient fractionation of the extracts shown in (A), with anti-γ-tubulin staining. Excess of GCPs 4+5 had no effect on the profile of the gradient, compared to control (excess of GCP4 only), whereas excess of GCPs 4+6 led to the formation of higher order complexes. Excess of GCPs 4+5+6 resulted into the complete sequestration of γTuSCs, and incorporation into γTuRCs. (C-E) Sucrose gradient fractionation of the extracts containing excess GCPs 4+5, 4+6 and 4+5+6, respectively. Fractions were separated by SDS-PAGE, blotted, and probed with antibodies against the different GCPs, in addition to γ-tubulin, to visualise the displacement of γTuSCs, and the proteins in excess (arrows). (F-H) Immunoprecipitations from fractions 2 to 7 of the gradients shown in C-E, using anti-GCP5 (F, H), or anti-GCP6 (G). The different immunoprecipitates were probed on Western blots with antibodies as indicated. (F) Excess GCP5 failed co-precipitate with significant amounts of other GCPs in fractions 2 to 5. (G) By contrast, excess GCP6 co-precipitated with GCPs 2, 3, 4, and γ-tubulin in intermediate-sized complexes in fractions 4 and 5. (H) Excess GCPs 4+5+6 resulted in the co-precipitation of the three proteins together with γ-tubulin (fraction 5).

Immunoprecipitation from the gradient fractions revealed that this shift was due to the binding of γTuSCs to assemblies of GCPs 4 and 6 (Fig. 3G). This is consistent with the idea of GCPs 4 and 6 forming an intermediate that can bind γTuSC, whereas GCP5 cannot efficiently bind to GCP4 in the absence of GCP6 (Fig. 3F). When an excess of all three GCPs 4, 5, 6 was created, immunoprecipitation of GCP5 yielded co-precipitating GCPs 4 and 6 (Fig. 3H), supporting the idea of a hierarchy of assembly, as proposed above, with GCPs 4/6 as the ultimate core. Interestingly, the high amounts of GCPs 4, 5 and 6 resulted in the disappearance of γTuSC-peaks, and led to an increase of γTuRCs (Fig. 3E). These results indicate that the GCP4/5/6 sub-complex binds γTuSCs and leads to their assembly into γTuRCs.

### Reconstituted γTuRCs from the GCP4/5/6 core are able to nucleate microtubules

To evaluate the capacity of the GCP4/5/6 sub-complex to promote the assembly of γTuRCs, we loaded the sub-complex onto beads and complemented these beads with γTuSCs, to assay for the presence and activity of reconstituted γTuRCs (Fig. 4A). In a first step, anti-GCP5 beads were used to immunoprecipitate either the γTuRC or the GCP4/5/6 sub-complex from a HEK293 cytoplasmic extract (“γTuRC extract”, prepared at 100mM or 600mM KCl respectively). We also used cells depleted of GCP2 to prepare extract at high salt, to ensure that the sub-complex was efficiently stripped of γTuSCs. The resulting GCP4/5/6-beads were washed and incubated at 100mM KCl with a second extract, prepared from cells depleted of GCPs 4, 5, and 6 (“γTuSC extract”, Fig. 4B). Reconstitution of γTuRCs was monitored by Western Blot (Fig. 4C), and by measuring the nucleation activity of the beads following incubation with pure tubulin (Fig. 4D-F). The GCP4/5/6-beads were lacking GCPs 2 and 3, compared to γTuRC beads, and they were not able to nucleate microtubules. Addition of the γTuSC extract resulted in re-incorporation of γTuSCs, and in recovery of the nucleation capacity of the beads. As a control, the γTuSC extract didn’t show any binding to the beads without GCP4/5/6, and these beads failed to nucleate any microtubules. Quantification of the number of microtubules nucleated per bead showed that reconstituted GCP4/5/6-beads with γTuSCs reached up to 78% of the nucleation capacity of the positive control, i.e. beads bound to native γTuRC. The difference in nucleation capacity compared to the positive control may reflect a decrease in the number of bound complexes (in particular after GCP2-depletion). Altogether, our results show that the GCP4/5/6 sub-complex can trigger the assembly of nucleation-competent γTuRCs.

**Figure 4:**
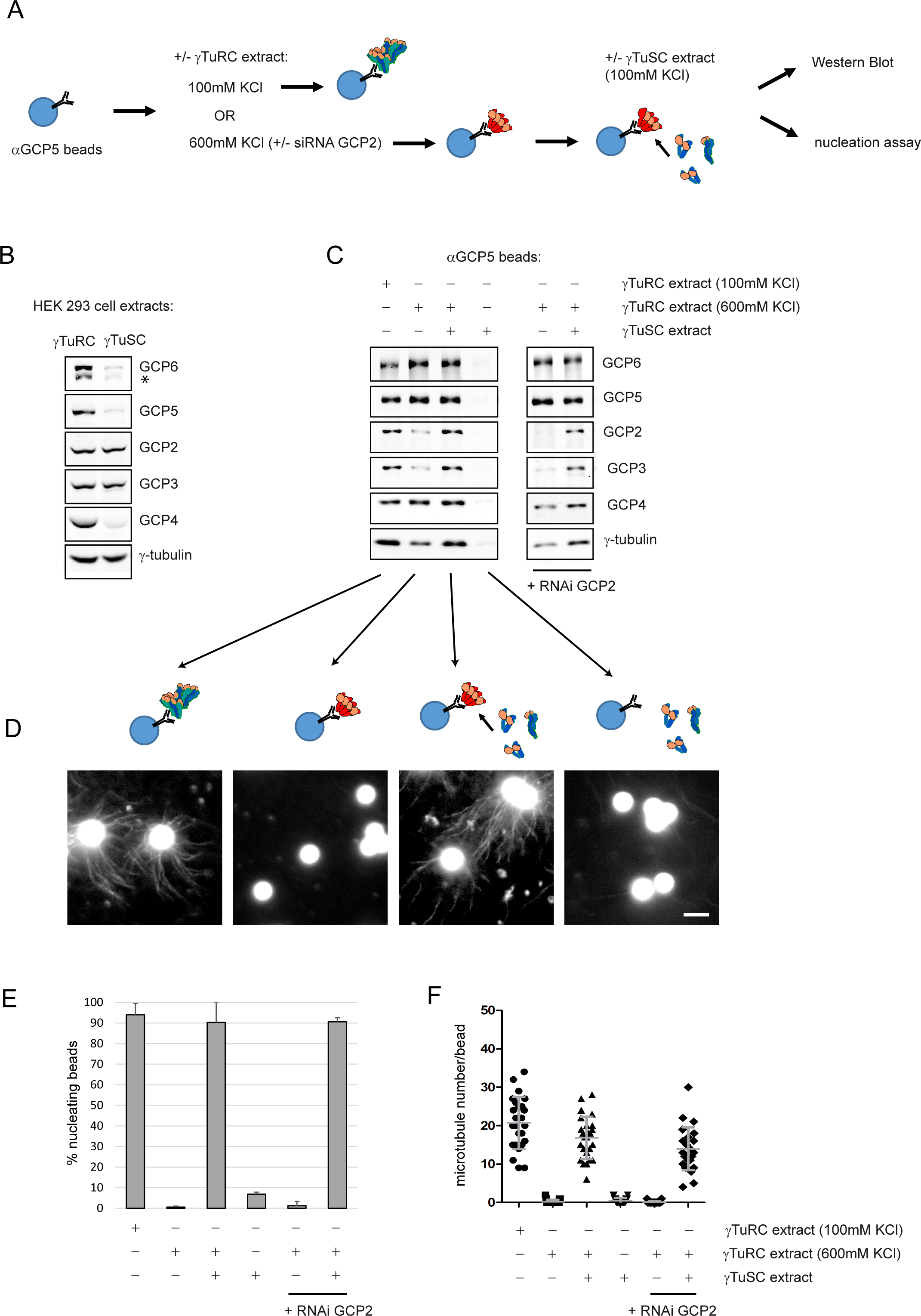
Reconstitution of functional γTuRCs from the purified GCP4/5/6 complex. (A) Experimental design. Dynabeads coupled with antibody against GCP5 were incubated with “γTuRC extract”: cytoplasmic extract from HEK293 cells lysed in the presence of 100 mM KCl, with or without adjustment to 600 mM KCl. Beads incubated with extracts at 600 mM KCl were subsequently rinsed and incubated with “γTuSC extract”: cytoplasmic extract from HEK293 cells treated with siRNAs against GCPs 4, 5, and 6, and lysed in the presence of 100 mM KCl. Following incubation, rinsed beads were used in a microtubule nucleation assay, or analysed on Western blots. (B) Western Blot analysis of the γTuRC and γTuSC extracts (*: degradation product of GCP6). (C) Left panel: Western Blot analysis of the eluates from the anti-GCP5 beads loaded with γTuRCs (γTuRC extract at 100 mM KCl), with the GCP4/5/6 complex (γTuRC extract at 600 mM KCl), with the GCP4/5/6 complex and in a second step with the γTuSC extract, or with the γTuSC extract alone. Right panel: anti-GCP5 beads loaded with the GCP4/5/6 complex prepared from GCP2 siRNA-treated cells, with or without addition of the γTuSC extract. (D) Microtubule nucleation from the anti-GCP5 beads loaded as in (C), incubated for 3 minutes at 37°C with pure TAMRA-labelled tubulin. Representative images are shown (beads are autofluorescent). Scale bar: 5μm (E) Percentage of beads showing radial microtubule arrays (“nucleating beads”; mean ± SD, 3 independent experiments, >100 beads counted / experiment). (F) Number of microtubules associated per bead (25 beads scored / condition, error bars: mean ± SD).

### Role of the GCP6-specific insertions in γTuRC assembly

Since GCP6 appears to play a central role in the assembly of the γTuRC, and since this protein is characterized by extensive non-grip sequences in its N-terminal region and in the domain connecting the grip1 and grip2 motifs (Fig. 5A, Fig. 6A), we constructed deletion mutants, to evaluate the importance of these sequences. The deletion mutants were expressed in stable HeLa cell lines in an inducible manner, following depletion of endogenous GCP6 by RNAi. We measured the potential of the mutants to rescue the assembly and function of γTuRCs by quantifying the amount of γTuRCs in sucrose gradients, and the formation of bipolar spindles in mitotic cells. In agreement with published data, depletion of GCP6 induces the loss of γTuRCs, the delocalisation of γ-tubulin from centrosomes and from the mitotic spindle, and inhibits the separation of the spindle poles (Fig. 5B, F; Bahtz et al., 2012; Farache et al, 2016; Cota et al., 2017). Deletion mutants were designed based on structure prediction. The wide central insertion (residues 675-1501) can be divided into three main regions. The first region (675-816) is predicted as a continuum of the helix α11 seen in GCP4, with a potential coiled-coil structure (residues 730-760). The middle region (816-1400) is unstructured, and contains a repeated sequence described to be phosphorylated by Plk4 (residues 1027-1269, Bahtz et al., 2012). The third region (1400-1501) is composed of multiple small helices (Fig. 5A). A combination of deletions of these domains was evaluated for rescue of spindle bipolarity (Fig. 5C-D). Depletion of GCP6 in wild-type cells resulted in more than 70% of mitotic cells with a monopolar spindle. The deletion mutants rescued the bipolarity of spindles, except if the third region (1400-1501) was absent. GCP6Δ675-1400 fully rescued spindle bipolarity, whereas GCP6Δ675-1501 showed no rescue at all (Fig. 5D). We observed that induction of GCP6 mutants with doxycycline leads to their overexpression, but the rescue was also observed in the absence of induction, due to a leak of the promoter (Fig. 5D-E). Sucrose gradient profiles confirm that γTuRCs were fully recovered when only the region [675-1400] was deleted (Fig. 5 F-G). These results show that most of the central insertion of GCP6 is not required for γTuRC assembly, except for the third region containing the last 100 amino acids. We immunoprecipitated the GCP4/5/6 complex at 500mM KCl, from cells expressing GCP6Δ675-1400 in low amounts (without addition of doxycycline). This deletion mutant of GCP6 still associated with GCP4 and GCP5, thus maintaining stable interactions at high salt (Fig. 5G).

**Figure 5:**
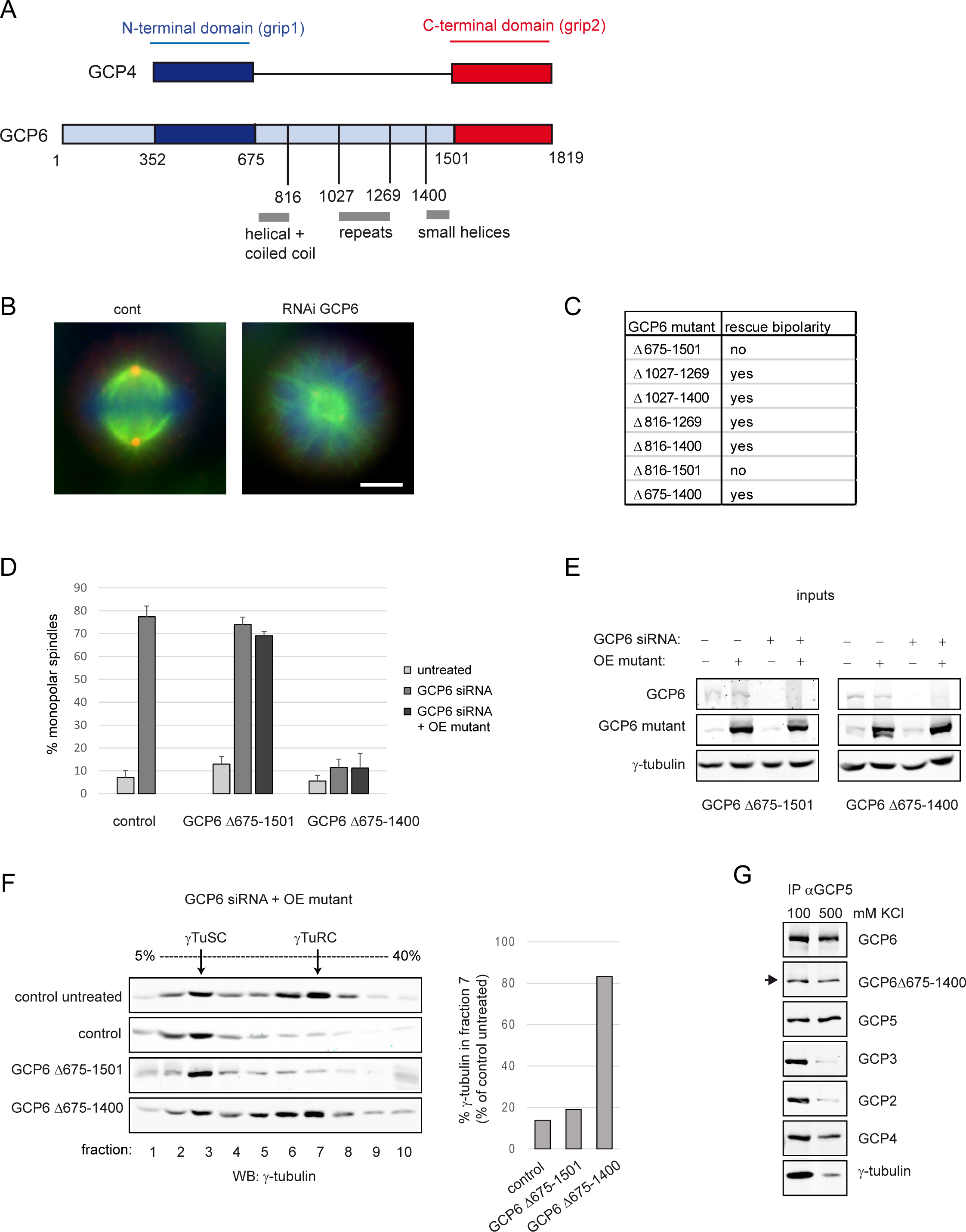
Structure/function of the GCP6-specific central insertion. (A) Schematic alignment of the primary structures of GCP4 and GCP6. The conserved N-terminal domain is boxed in blue, the C-terminal domain in red. The specific extension and insertion in GCP6 are in light blue. The positions of the GCP6 amino acids at the junctions between domains are indicated, together with the positions of the deletions. Predicted structural features are in grey. (B) Comparison of mitotic spindles in control and GCP6 siRNA-treated cells, with γ-tubulin stained in red, microtubules in green, and DNA in blue. Scale bar, 5 μm. (C) Summary of the functionality of the GCP6 deletion mutants, to rescue spindle bipolarity in mitotic cells. Deletion mutants were expressed from stably transfected cells, with or without induction, after treatment with GCP6 siRNA. The constructs were resistant to the siRNA. (D) Percentage of monopolar spindles in control cells and in cell lines expressing GCP6Δ675-1501 or GCP6Δ675-1400, with or without GCP6 siRNA treatment, with or without overexpression (OE) following induction by doxycycline. GCP6Δ675-1400 rescues spindle bipolarity, even without induction, due to leaky expression (mean ± SD, 3 independent experiments, 100 cells counted / experiment). (E) Western Blot analysis of cytoplasmic extracts from cell lines expressing GCP6Δ675-1501 or GCP6Δ675-1400, with or without GCP6 siRNA treatment, with or without overexpression (inputs for F-G). Blots were probed with antibodies against GCP6 (top and middle rows), or antibody against γ-tubulin (bottom rows). (F) Sucrose gradient fractionation of extracts from control cells or cell lines expressing GCP6Δ675-1501 or GCP6Δ675-1400, after GCP6 siRNA treatment and overexpression of the mutants. The top gel shows an untreated control (no siRNA). Fractions were separated by SDS-PAGE and analysed on Western Blots, probed with antibody against γ-tubulin. The graph on the right shows the quantification of γ-tubulin signal in fraction 7 of the gels displayed, relative to the untreated control cells. (G) Immunoprecipitation of GCP5 from a cytoplasmic extract prepared from the cell line expressing GCP6Δ675-1400, without overexpression, at 100 mM or 500 mM KCl. Co-precipitation of the mutant (arrow) is similar at both concentrations.

**Figure 6:**
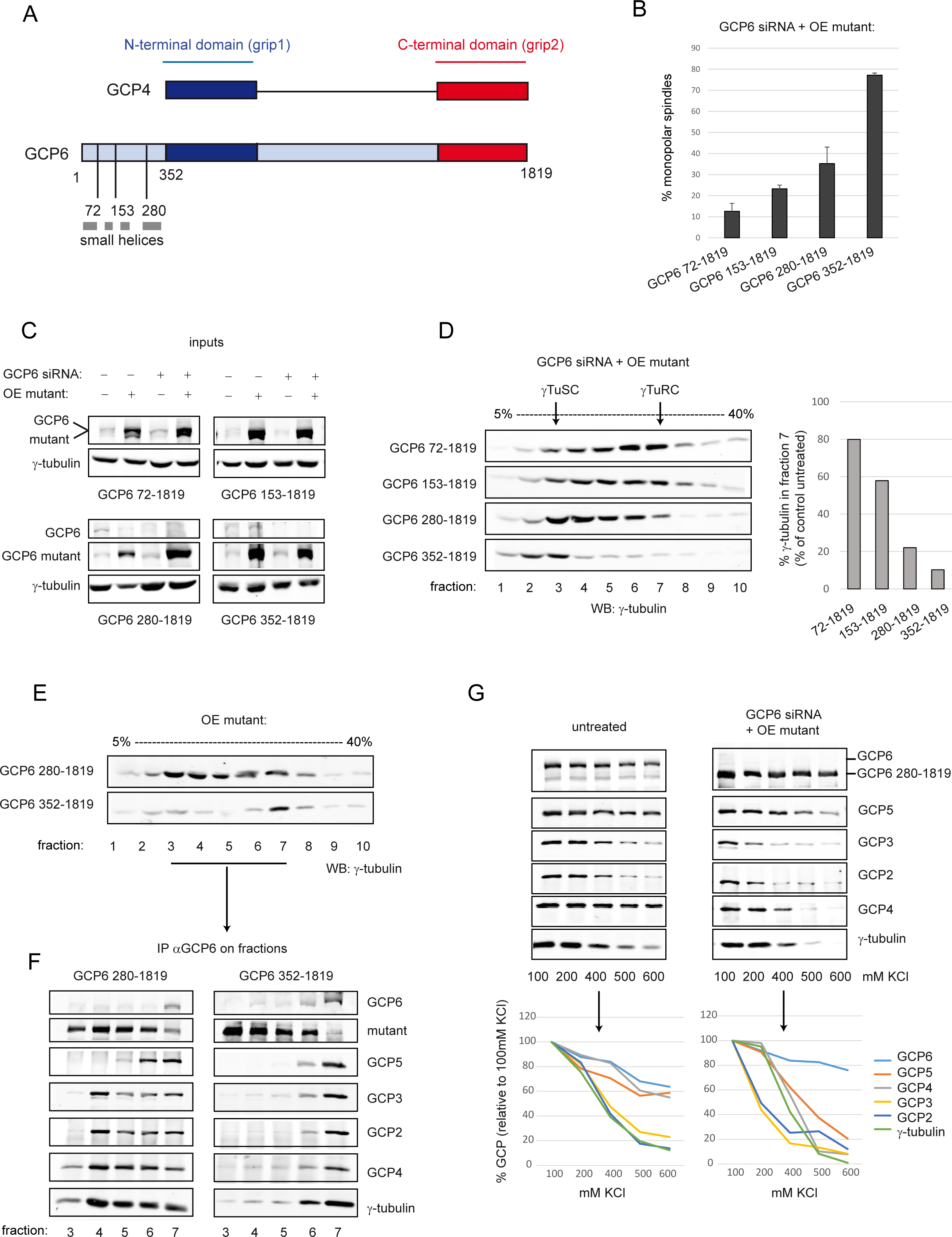
Structure/function of the GCP6-specific N-terminal extension. (A) Schematic alignment of the primary structures of GCP4 and GCP6, with the positions of the deletions indicated in the N-terminal extension of GCP6. Predicted helical regions are indicated in grey. (B) Percentage of monopolar spindles in stable cell lines expressing different GCP6 deletion mutants, after siRNA treatment and overexpression of the mutants (mean ± SD, 3 independent experiments, 100 cells counted / experiment). (C) Western Blot analysis of cytoplasmic extracts from the stable cell lines, with or without GCP6 siRNA treatment, with or without overexpression of the mutants (inputs for D-G). The blots were probed with antibodies against GCP6 (top rows), or γ-tubulin (bottom rows). (D) Sucrose gradient fractionation of extracts from the cell lines, after GCP6 siRNA treatment and overexpression of the mutants. The graph on the right shows the quantification of γ-tubulin signal in fraction 7 of the blots displayed, relative to the untreated control shown in Fig 5F. (E) Sucrose gradient fractionation of the extracts from cell lines overexpressing GCP6 280-1819 and GCP6 352-1819. Western blots were probed with antibody against γ-tubulin. (F) Immunoprecipitation of the overexpressed mutants from fractions 3 to 7 of the gradients shown in (E). GCP6 280-1819 co-precipitates with GCPs 2, 3, 4 and γ-tubulin (fractions 4, 5), but GCP6 352-1819 shows no interaction. (G) Immunoprecipitation of GCP6 280-1819 from extracts of untreated control cells (left panel), or cells treated with GCP6 siRNA, and induced to overexpress of the mutant (right panel). Extracts were supplemented with increasing concentrations of KCl (100 to 600 mM). The graphs below represent the percentage of co-immunoprecipitated GCPs at the different KCl concentrations, quantified from the intensities of the bands and relative to the amounts precipitated at 100 mM KCl.

To investigate the role of the N-terminal extension of GCP6, it was gradually shortened by cutting between its multiple predicted helices (Fig. 6A). Deletion of the entire extension (GCP6 352-1819) abolished the capacity of GCP6 to rescue spindle bipolarity, and prevented the assembly of γTuRCs. All other mutants rescued to various extents, but the efficiency of the rescue gradually decreased with the length of the deletion (Fig. 6B-D; suppl. Fig. 6). GCP6 280-1819 generated low amounts of γTuRCs, and complexes of intermediate size accumulated on sucrose gradients. These complexes may represent partially assembled γTuRCs, or unstable γTuRCs that started to disassemble during sedimentation. To evaluate if the GCP6 mutants 280-1819 or 352-1819 affected the interactions with other GCPs of the γTuRC, we overexpressed them and performed immunoprecipitations from gradient fractions, in an equivalent manner as in Fig 3. GCP6 280-1819 co-immunoprecipitated with GCP4 and with γTuSCs, to similar degrees as the wild-type protein, showing that the protein interactions were maintained. By contrast, GCP6 352-1819 failed to interact with the other GCPs (Fig. 6E-F). We then immunoprecipitated GCP6 280-1819 from cytoplasmic extracts at increasing salt concentrations. Although GCPs 2, 3, 4, 5 efficiently co-precipitated at 100mM KCl, their interaction was lost at 500mM KCl. At 200mM KCl, GCP2 and GCP3 still interacted with wild-type GCP6, but their interaction with the 280-1819 mutant was strongly reduced (Fig. 6G). Altogether, these results suggest that the region 280-352 of GCP6 is necessary for the interactions with GCP4 and with γTuSCs, and that the first 280 residues of GCP6 stabilise its interactions with the γTuRC.

## Discussion

We demonstrate that GCPs 4, 5, and 6 form a stable sub-complex that permits the association with γTuSCs into a functional γTuRC. Within the sub-complex, the stoichiometric ratio of these GCPs matches the values found in recently reported structures of native γTuRCs (Wieczorek et al., 2020; Consolati et al., 2019; Liu et al., 2019). We find that GCPs 2 and 3 are also present in our preparations of the sub-complex, but at a molar ratio equivalent to half-a-γTuSC. We believe that this reflects stochastic binding of remnant γTuSCs after incomplete salt-stripping at 500mM KCl. More stringent conditions with even higher salt concentrations failed to increase the purity of the sub-complexes, but rather led to their disassembly (Fig. 1A).

### Role of the GCP4/5/6 sub-complex in the assembly and stabilization of the γTuRC

Structural and biochemical work has shown that γ-tubulin complexes in budding yeast are formed by lateral assembly of γTuSCs (Kollman et al., 2010). Lateral association and oligomerization into a helically shaped template are supported by targeting factors, such as Spc110, that promote the recruitment of γTuSCs to the spindle pole body and thus couple localization to the assembly into a nucleation-competent complex (Kollman et al., 2010; Lyon et al., 2016; Lin et al., 2014; 2016). Other targeting factors have been identified in different species of yeast, including Spc72, Mozart1, or Mto1/2 (Lin et al., 2016; Masuda and Toda, 2016; Lynch et al., 2014; Leong et al., 2019). In most eukaryotes, fully assembled, helically shaped complexes are already present as soluble entities in the cytoplasm, in the form of γTuRCs. Nevertheless, these soluble γTuRCs remain inactive unless recruited to specific sites of microtubule nucleation, thus allowing tight spatial and temporal control of microtubule formation (Farache et al., 2018). GCPs 4, 5, and 6 are essential for the assembly and/or for the stabilization of γTuRCs, since depletion of either component causes a reduction of γTuRCs both at the centrosome and in the cytoplasm (Izumi et al., 2008; Bahtz et al., 2012; Scheidecker et al., 2015; Farache et al., 2016; Cota et al., 2017). Besides GCPs 4, 5, 6, additional factors may still be needed, such as Mozart1, actin, or other proteins corresponding to unassigned densities in cryo-electron microscopy structures of native γTuRCs (Lin et al., 2016; Cota et al., 2017; Wieczorek et al., 2020; Consolati et al., 2019; Liu et al., 2019). It is now clear that GCPs 4, 5, and 6 integrate into the helical wall of the γTuRC, and that they are laterally bound to γTuSCs, but their specific role within the γTuRC still remains to be determined (Wieczorek et al., 2020; Consolati et al., 2019; Liu et al., 2019). Our results show that they form a nucleus promoting the stable assembly of the 2MDa complex. During the formation of γTuRCs, complexes of GCPs 4, 5, 6 may act as building blocks that recruit γTuSCs by lateral association and thereby initiate γTuRC-assembly. This hypothesis is directly supported by our experiments with salt-stripped sub-complexes of GCPs 4, 5, 6 that can drive the formation of functional γTuRCs, after incubation with γTuSC-containing cytoplasm (Fig. 4).

The assembly of γTuRCs via GCPs 4, 5, 6 may be important for the regulation of its microtubule-nucleation activity: cryo-electron microscopy structures show an asymmetric architecture, incompatible with the geometry of microtubules (Wieczorek et al., 2020; Consolati et al., 2019; Liu et al., 2019). The presence of GCPs 4, 5, and 6 may therefore prevent soluble γTuRCs from acquiring an active conformation, unless bound to additional activating factors, such as Cdk5rap2/Cep215, at designated microtubule-organizing centres. Consistently, the GCP5 subunit was identified in two different conformations at position 10 within the γTuRC, and conformational changes propagated towards positions 11 to 14 of the γTuRC-helix, suggesting that these might regulate the overall structure and the activation of the γTuRC (Wieczorek et al., 2020). In addition, specific non-grip domains in GCP6 might also be involved in the binding to regulatory factors. The large central insertion in GCP6, including nine tandem repeats of 27 amino acids, has been proposed to be regulated by Plk4-dependent phosphorylation (Bahtz et al., 2012). Contrary to this view, our deletion experiments show that the majority of the central insertion (residues 675-1400, comprising the tandem repeats) can be removed from GCP6, without impacting the assembly or the activity of the γTuRC, since a Δ675-1400 deletion mutant permits the assembly of regular mitotic spindles. Moreover, neither the recruitment of γTuRCs to the centrosome nor to the mitotic spindle were affected by this deletion.

Besides any regulatory role, the GCP4/5/6 sub-complex may contribute to the stabilization of γTuRCs post-assembly, by preventing the loss of γTuSCs from the helical complex. In particular, the N-terminal extension of GCP5 may be part of a lumenal bridge within the γTuRC and may thereby fulfil a stabilizing role (Wieczorek et al., 2020). Moreover, the N-terminal extension of GCP6 (amino acids 1-352) may correspond at least in part to a stabilizing “plug” seen in the cryo-electron microscopy structure of the γTuRC (Wieczorek et al., 2020). In accordance, we have observed that progressive shortening of the N-terminal extension of GCP6 destabilises the γTuRC and renders it more sensitive to salt treatment. For example, the deletion of the first 279 amino acids leads to the loss of γTuSCs at 200mM KCl, and to the loss of GCPs 4 and 5 at 400mM KCl, whereas wild type γTuRCs remain stable under these conditions (Fig. 6G). Additional stabilization of the γTuRC may be provided by contacts between the central insertion in GCP6 (amino acids 675-1501) and GCP2 in position 13, since GCP6 deletion mutants that lack this insertion (Δ675-1501) prevent the formation of stable γTuRCs (Fig. 5F; Wieczorek et al., 2020).

Overall, the sub-complex of GCPs 4, 5, and 6 may promote lateral associations with γTuSCs via the grip1 domains of its peripheral constituents GCP4 and GCP6, and may stabilize interactions within a larger γTuRC via the N-terminal extensions of GCPs 5 and 6.

The question has been raised whether γ-tubulin and GCP4 assemble with either GCP5 or GCP6 into intermediate hetero-tetramers, such as “γ-TuG4/5” and “γ-TuG4/6”, equivalent to γTuSCs (Wieczorek et al., 2020; Liu et al., 2019). Our data don’t provide direct evidence for this mechanism, since salt extraction or GCP2 depletion yields “monolithic” sub-complexes in which GCPs 4, 5, and 6 are stably bound to each other. In fact, salt extraction demonstrates that the affinities between the GCPs of this sub-complex are higher than the affinities between γTuSCs.

Nevertheless, we have seen that “γ-TuG4/6” intermediates form in the absence of GCP5, and that these are sufficient to establish lateral contacts with γTuSCs, since excess protein levels of GCPs 4 and 6 are able to co-immunoprecipitate with GCPs 2 and 3, without GCP5 (lanes 4, 5 in Fig. 3G). By contrast, GCP5 needs the presence of both GCPs 4 and 6 in order to form any higher-order complex, suggesting that in human cells, a “γ-TuG4/5” does not exist (Fig. 2). The situation may be different in other species, such as fission yeast, where the orthologs of GCPs 4 and 5, Gfh1p and Mod21p, can be co-immunoprecipitated in the absence of the GCP6 ortholog Alp16 (Anders et al., 2006). In both experimental systems, though, the GCP6 ortholog needs GCP4 to associate with GCP5, suggesting that the spatial arrangement of GCPs 4, 5, and 6 within γ-tubulin complexes may be conserved across species. Based on our experiments using depletion or overexpression of individual GCPs (Figs. 2, 3), we propose a hierarchy of assembly, with GCPs 4 and 6 together enabling the recruitment of GCP5 into a stable sub-complex that drives lateral association with γTuSCs into a full-sized γTuRC (Fig. 7). Studies by Cota et al. (2017) and Farache et al. (2016) show that depletion of either GCP4 or GCP5 leads to milder defects than the depletion of GCP6, confirming a central role of GCP6 in γTuRC-assembly. This raises the possibility that partial or unstable γ-tubulin complexes may still form by lateral interactions between GCP6 and γTuSCs, or that γTuSCs assemble into larger complexes *in situ*, at centrosomes or spindle pole bodies, partially stabilized by GCP6, and with restricted nucleation capacity (Vérollet et al., 2006; Xiong and Oakley, 2009; Masuda and Toda, 2016).

**Figure 7:**
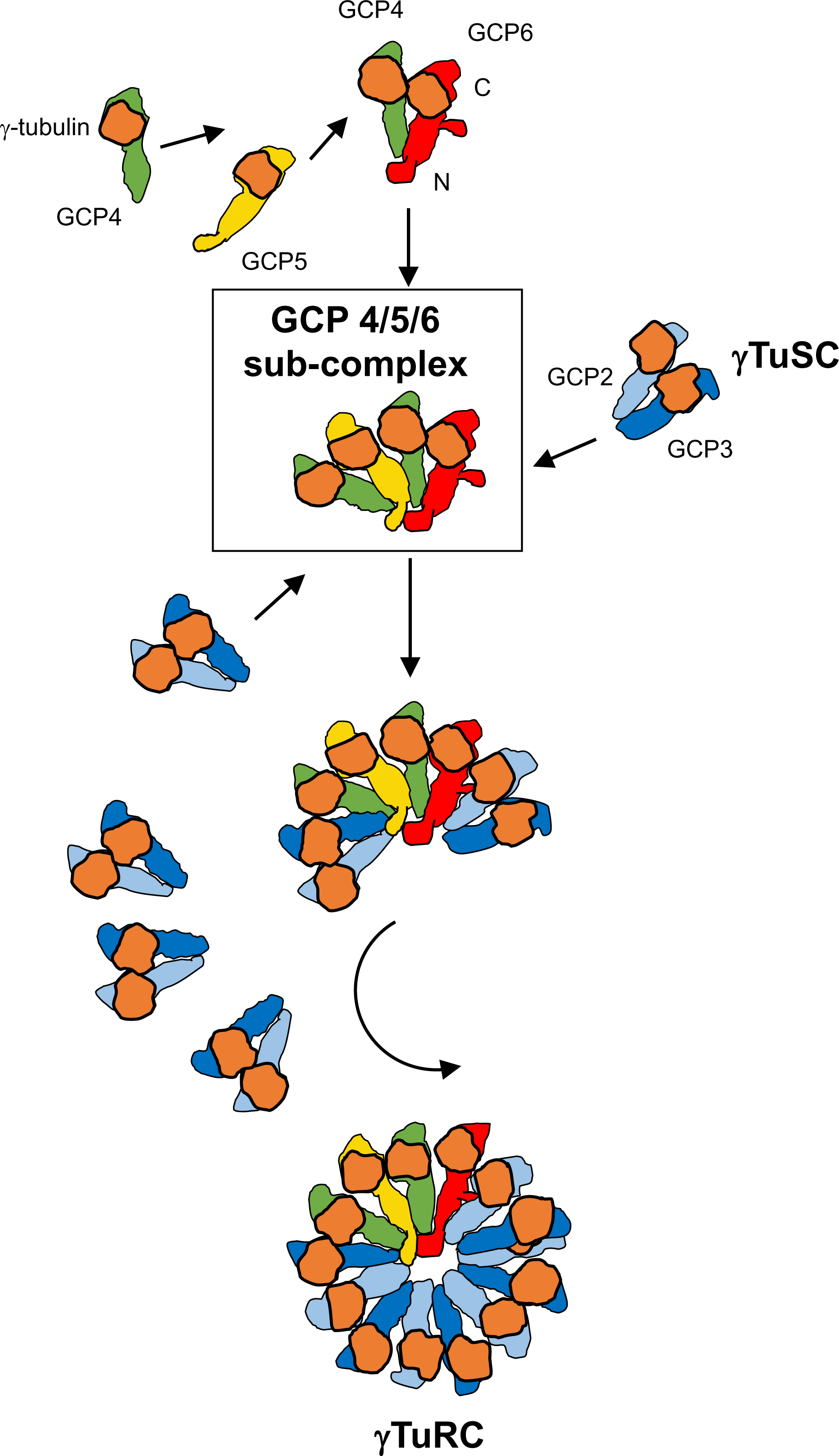
Model of γTuRC assembly. We propose that lateral alignment of GCP6 with GCP4 enables the binding of GCP5 and an additional copy of GCP4, to form a stable intermediary, the GCP 4/5/6 sub-complex. The sub-complex can then drive and stabilize the association with γTuSCs, until one complete helical turn is reached. Sequence extensions in the N-terminal regions of GCPs 5 and 6, as well as an insertion between the grip1 and grip2 motifs of GCP6 contribute to the stabilization of interactions with neighbouring γTuSCs, and across the lumen of the γTuRC.

For the future, to gain further insights into the assembly of γTuRCs, it would be interesting to perform controlled reconstitution of the multiprotein complex from purified, recombinant components.

## Materials and Methods

### Cell culture and generation of stable cell lines

HeLa Flp-In T-REx (Tighe et al., 2008), and HEK293 FT cells were grown at 37°C, at 5% CO_2_ in Dulbecco’s modified Eagle’s medium (DMEM), supplemented with 10% fetal bovine serum.

For CRISPR/Cas9 targeting of GCP6 in HEK293 FT cells, a target guide RNA overlapping the stop codon (5’-AACTACTACCAGGACGCCTG-3’, computed from http://crispr.mit.edu/) was inserted into pSpCas9(BB)-2A-Puro (Addgene). The homologous recombination donor was a DNA fragment consisting of a 1445 bp left homology arm (GCP6 exon 20 to 25), a 2244 bp insertion sequence (GST-6His coding sequence, in frame with exon 25 of GCP6, followed by a puromycin resistance cassette), and a 1577 bp right homology arm (GCP6 exon 25 and 3’ untranslated sequence). Cells were transfected using Lipofectamine 2000 (Invitrogen, Carlsbad, CA). Limiting dilution cloning was performed 3 days post-transfection, in conditioned medium supplemented with 2 μg/ml puromycin. Targeted clones were identified by PCR, and verified by sequence analysis and by Western Blotting, using an anti-GST antibody (Roche Applied Science, Basel, CH).

HeLa Flp-In T-REx overexpressing cell lines were obtained as described (Farache et al., 2016). GCP5 and GCP6 (resistant to siRNA) were expressed from pCDNA5/FRT/TO (Invitrogen, Carlsbad, CA). Internal deletions of GCP6 were constructed using the Gibson kit (NEB, Ipswich, MA), whereas N-terminal deletions were generated by PCR (using restriction sites BamH1 and Ale1 in the sequences of the vector or the GCP6 DNA, respectively). Briefly, GCP6 constructs were co-transfected with pOG44 (Invitrogen, Carlsbad, CA) expressing the Flp recombinase, using CaCl_2_. Resistant clones at 200μg/ml hygromycin B were picked and expanded to obtain clonal cell lines. The phenotypes of at least two independent clones were compared for each construct. Transient transfection of the GCP5 construct was performed using Lipofectamine 2000. Transgene expression was induced using 1μg/ml doxycycline. For RNA interference, 10 nM siRNA were transfected into HeLa Flp-In T-REx cells, using Lipofectamine RNAi max (Invitrogen, Carlsbad, CA; Farache et al., 2016). The medium was replaced after 24 h, with or without doxycycline. Cells were harvested or fixed at 72 h post-transfection.

### Purification of the GCP4/5/6 sub-complex

Cells homozygous for GCP6-GST-6His were harvested by trypsin-treatment, rinsed with PBS and stored at −80°C. Pellets were resuspended in 50 mM HEPES/KOH, at pH 7.2, containing 1 mM MgCl_2_, 1 mM EGTA, and 100 mM KCl (HB100), and supplemented with 0.1 mM GTP, 1 mM DTT, 1 mM PMSF, and complete protease inhibitor cocktail (Roche, Basel, CH). Cells were disrupted by sonication and centrifuged 30 min at 30,000 g. Protein precipitation from the supernatant was performed by adding 30% of a saturated (NH_4_)_2_SO_4_ solution, followed by solubilization in HB500 (HB with 500mM KCl), supplemented with 1 mM DTT, 1 mM PMSF, protease inhibitors, 15 mM imidazole and 0.05 % Igepal CA-630. The solution was added to glutathione Sepharose 4B (GE Healthcare, Chicago, IL) and incubated 4 h at 4°C under agitation. Beads were washed twice with binding buffer, and twice with HB500, supplemented with 1 mM DTT, 15 mM imidazole and 0.05% Igepal CA-630. Elution was performed with the same buffer containing 40 mM reduced glutathione, pH 7.2. Buffer was exchanged by passing the eluate through a desalting PD-10 column (GE Healthcare, Chicago, IL), equilibrated with 50 mM HEPES/KOH, pH 7.2, containing 1 mM MgCl_2_, 500 mM KCl, 0.5 mM DTT and 0.05% Igepal CA-630. The solution was then incubated overnight with Ni-NTA agarose (Qiagen, Hilden, DE) at 4°C, and washed twice with binding buffer, and twice with the same buffer supplemented with 15 mM imidazole. Elution was performed with 200 mM imidazole. Eluted proteins were analyzed on Western blots, or on polyacrylamide gels with the PlusOne silver staining kit (GE Healthcare, Chicago, IL). Size exclusion chromatography was performed on Superdex™ 200 Increase 5/150 GL (GE Healthcare, Chicago, IL).

### Sucrose gradient sedimentation

Sucrose gradients were performed as described in Farache et al. (2016), in HB100, or at varying concentrations of KCl.

### Immunoprecipitation

Cytoplasmic lysates were produced from trypsinized HeLa Flp-In T-Rex cells, lysed in HB100 supplemented with 1 mM GTP, 1 mM DTT, 1 mM PMSF, protease inhibitors, 1% Igepal CA-630 and 10% glycerol. After 5 minutes on ice, cells were centrifuged for 10 minutes at 16,000 g. Aliquots of the supernatant containing 500 μg of protein were diluted in 100 μl buffer. These cytoplasmic lysates where then supplemented with KCl, to increase the concentration as needed, before incubation with the anti-GCP4, anti-GCP5, or anti-GCP6 antibodies, for 2h at 4°C. 50 μl of protein A-dynabeads (Invitrogen, Carlsbad, CA) were added for 1h at 4°C, and proteins were eluted in gel loading buffer, after two washes in HB100 containing 10% glycerol.

Immunoprecipitation from the gradient fractions was performed as above, by adding antibodies to the fractions and incubating for 2 h at 4°C, followed by incubation with the protein A-dynabeads for an additional hour. The beads were then washed twice in HB100 containing 10% glycerol, and samples were eluted in gel loading buffer.

### Microtubule nucleation from beads

Dynabeads were washed and incubated with the anti-GCP5 antibody in PBS, containing 0.02% Tween-20, for 2.5h at 4°C. After three additional washes, HEK293 FT cytoplasmic extracts (“γTuRC extracts”) were added to the beads and incubated for 2h at 4°C. Extracts were prepared as described in the immunoprecipitation protocol, at 100 or 600 mM KCl. For each reaction, we used cells from one dish of 100 mm diameter (yielding 4 mg of protein), lysed in 100 μl buffer, for 5 μl beads.

After three washes with HB100 + 10% glycerol, a freshly prepared cytoplasmic extract depleted of GCPs 4, 5, and 6 (“γTuSC extract”) was added to the beads. Again, cells from one dish of 100mm diameter were lysed in 100 μl buffer, at 100 mM KCl. After 2h incubation at 4°C, beads were washed three times with BRB80 (80 mM PIPES, pH 6.9, 1 mM EGTA, 1 mM MgCl_2_), and resuspended in 12.5 μl BRB80.

Nucleation was tested by incubating 2.5 μl beads for 3 minutes at 37°C with a solution containing 1 μl tubulin at 10mg/ml, 1 μl TAMRA-labeled tubulin at 2mg/ml, and 0.5 μl 10 mM GTP. The reaction was stopped by adding 45 μl of 1% glutaraldehyde in BRB80, for 5 minutes at 37°C. The beads were diluted in BRB80 and layered onto a cushion of 10% glycerol in BRB80, centrifuged onto coverslips and mounted in Vectashield solution (Vector Laboratories, Peterborough, UK).

Proteins were eluted from the remaining beads in gel loading buffer, and analyzed by Western Blotting.

### Western Blot analysis

Proteins were detected using an Odyssey imaging system (Li-cor Biosciences, Lincoln, NE), according to the manufacturer’s protocol, with IRDye 800CW- and 680CW-conjugated secondary antibodies (Invitrogen, Carlsbad, CA). Protein levels were quantified using Odyssey 2.1 software. The Odyssey fluorescence system provides a linear relationship between signal intensity and antigen loading. Band intensities were measured after background subtraction. For the quantification of γ-tubulin in fractions of sucrose gradients, the band intensities of individual fractions were normalized to the total amount of γ-tubulin in the experiment, corresponding to the sum of the intensities of all fractions of the sucrose gradient.

### Fluorescence microscopy

Cells grown on coverslips were fixed in methanol at −20°C, and processed for immunofluorescence, following standard protocols. Fluorescence microscopy was performed on a wide-field microscope (Axiovert 200 M; Carl Zeiss, Oberkochen, D) equipped with a Z motor, using a 63X (Plan Apo, 1.4 NA) objective. Images were acquired with an MRm camera and Axiovision software. Image processing was performed using Adobe Photoshop.

### Antibodies

Rabbit anti-GCP6 (Abcam, Cambridge, UK); mouse anti-γ-tubulin TU-30 (Exbio, Vestec, CZ); rabbit anti-GCP4 (Fava et al., 1999); rabbit anti-GCP2 (Haren et al., 2006); rabbit anti-γ-tubulin R75 (Julian et al., 1993); mouse anti-GCP3 C3, mouse anti-GCP5 E1, rabbit anti-GCP5 H300, mouse anti-GCP6 H9 (Santa Cruz Biotechnology, Santa Cruz, CA); mouse anti-α-tubulin (Sigma Aldrich, St. Louis, MO).

## Acknowledgements

We thank Valérie Guillet (Toulouse) for help with size exclusion chromatography, Stephen Taylor (Manchester) for the gift of HeLa Flp-in T-Rex, Nathalie Morin (Montpellier) for the gift of TAMRA-tubulin, and Jens Lüders (Barcelona) for fruitful discussions during the course of this work. We thank all our colleagues at the Université Paul Sabatier for critical discussion and for technical help. The project was in part supported by grant 13-BSV8-0007-01 from “Agence Nationale de la Recherche”, and by grant SFI20121205511 from “Fondation ARC pour la recherche sur le cancer”.

No competing interests declared.

**<u>Supplement to Figure 1:</u>** Western Blot analysis of sucrose gradient fractionation of HeLa cytoplasmic extract at different KCl concentrations. (A) The extract was prepared and centrifuged at 100 mM KCl. (B) The extract was adjusted to 500 mM KCl and centrifuged at the same concentration. (C and D) The extract was adjusted to 500 mM KCl and centrifuged at 200 and 100 mM KCl, respectively. All GCPs were probed in addition to γ-tubulin. Note the high levels of GCP4, spread in fractions 2 to 7 in (A).

**<u>Supplement to Figure 2:</u>** Western Blot analysis of sucrose gradient fractionation of cytoplasmic extracts from cells treated with siRNAs against GCPs 2, 4, 5, or 6.

**<u>Supplement to Figure 6:</u>** (A) Western Blot analysis of sucrose gradient fractionation of cytoplasmic cell extracts from the cell lines expressing GCP6 72-1819, GCP6 153-1819, GCP6 280-1819 or GCP6 352-1819, after GCP6 siRNA treatment and overexpression of the mutants. (B) Western Blot analysis of sucrose gradient fractionation of cytoplasmic cell extracts from the cell lines expressing GCP6 280-1819 or GCP6 352-1819, after overexpression of the mutants, without siRNA treatment.

